# Enrichment of disease-associated genes in cortical areas defined by transcriptome-based parcellation

**DOI:** 10.1101/2020.03.02.971911

**Authors:** Gryglewski Gregor, Murgaš Matej, Michenthaler Paul, Klöbl Manfred, Reed Murray Bruce, Unterholzner Jakob, Lanzenberger Rupert

## Abstract

The parcellation of the cerebral cortex serves the investigation of the emergence of uniquely human brain functions and disorders. We employed hierarchical clustering based on comprehensive transcriptomic data of the human cortex in order to delineate areas with distinct gene expression profiles. These profiles were analyzed for the enrichment of gene sets associated with brain disorders by genome-wide studies (GWAS) and expert curation. This suggested new roles of specific cortical areas in psychiatric, neurodegenerative, congenital and other neurological disorders while reproducing some well-established links for movement disorders and dementias. GWAS-derived gene sets for psychiatric disorders exhibited similar enrichment patterns in the posterior fusiform gyrus and inferior parietal lobule driven by pleiotropic genes. This implies that the effects of risk variants shared between neuropsychiatric disorders might converge in these areas. For several diseases, specific genes were highlighted, which may aid the discovery of novel disease mechanisms and urgently needed treatments.

## Introduction

The evolution of brain functions which underlie several uniquely human behaviors was paralleled by the specialization of the cerebral cortex.^1^ Accordingly, cortical parcellation was among the earliest pursuits in modern neuroscience. It has continued to advance our understanding by providing definitions of discrete areas and networks which facilitate specific scrutiny, interpretation of results and communication between researchers.^2^ Departing from the approach by Korbinian Brodmann based on cytoarchitectonic features assessed in a single specimen,^3^ methods which integrate a spectrum of different properties are continuously being developed with increasing standards pertaining to standardization and replicability. The influence of cortical parcellation is particularly pronounced in the brain imaging field where popular reference atlases have been applied in thousands of studies.^4^ Parcellations based on imaging outcomes assessed *in vivo*, e.g. brain structure, activation during tasks or at rest, structural or functional connectivity have been developed.^5^ While their replicability and utility for specific analyses could be demonstrated, their exact neurobiological underpinnings need to be established.^6^ Meanwhile, a large number of well-defined tissue properties can be assessed in cortical tissue *post mortem*.^7^ By performing automated analyses of cytoarchitecture on multiple brains, probabilistic atlases of cortical areas could be created.^8^ Interestingly, the borders of these areas were found to be aligned with gradients in multi-receptor density profiles measured using autoradiography,^9^ which inspired parcellation based on *in vivo* molecular imaging data.^10,11^ Despite these promising findings, the number of targets measurable by molecular imaging is highly limited and therefore may be insufficient to detect aspects of cortical differentiation which might become evident with a more extensive characterization of molecular tissue properties.

Therefore, the current work capitalizes on the human brain transcriptome data provided in the scope of the Allen Human Brain Atlas (AHBA) project.^12^ Currently, this data constitutes the most comprehensive characterization of gene expression in the human cerebral cortex. We have previously developed a method to generate continuous and unbiased estimates of gene expression in the entire cortex from discrete microarray sample data.^13^ This enabled the transcriptome-based parcellation of the cerebral cortex presented here. We performed hierarchical clustering based on data on the expression of 18,686 genes throughout the cerebral cortex. We expected that the distinct transcriptomic profiles of the cortical areas delineated in this manner would enable insights on their role in specific brain functions and vulnerability to disease. To this end, we analyzed these profiles for the enrichment of genes associated with specific brain disorders or related phenotypes by genome-wide studies (GWAS) and expert curation.^14^ Next to the discovery of novel associations between cortical areas and disorders, we calculated the relative contribution of genes to these associations to aid the prioritization of molecular targets for the development of urgently needed treatments.

## Results

### Transcriptome based parcellation of the human cerebral cortex

Hierarchical clustering of the cerebral cortex into 100 clusters and the corresponding dendrogram are shown in Figure 1. Transcriptome-based parcellation is presented alongside other established parcellations based on cytoarchitecture, functional and structural brain imaging and heritability of structural features in Supplementary Figure S3. Despite the high overlap observed for certain areas in each atlas, the adjusted Rand index (ARI), which reflects the overall similarity of two clusterings, ranged from 0.11 to 0.33. Notably, the maximum ARI observed between the different published parcellations was 0.37, indicating low similarity.

**Figure 1.**
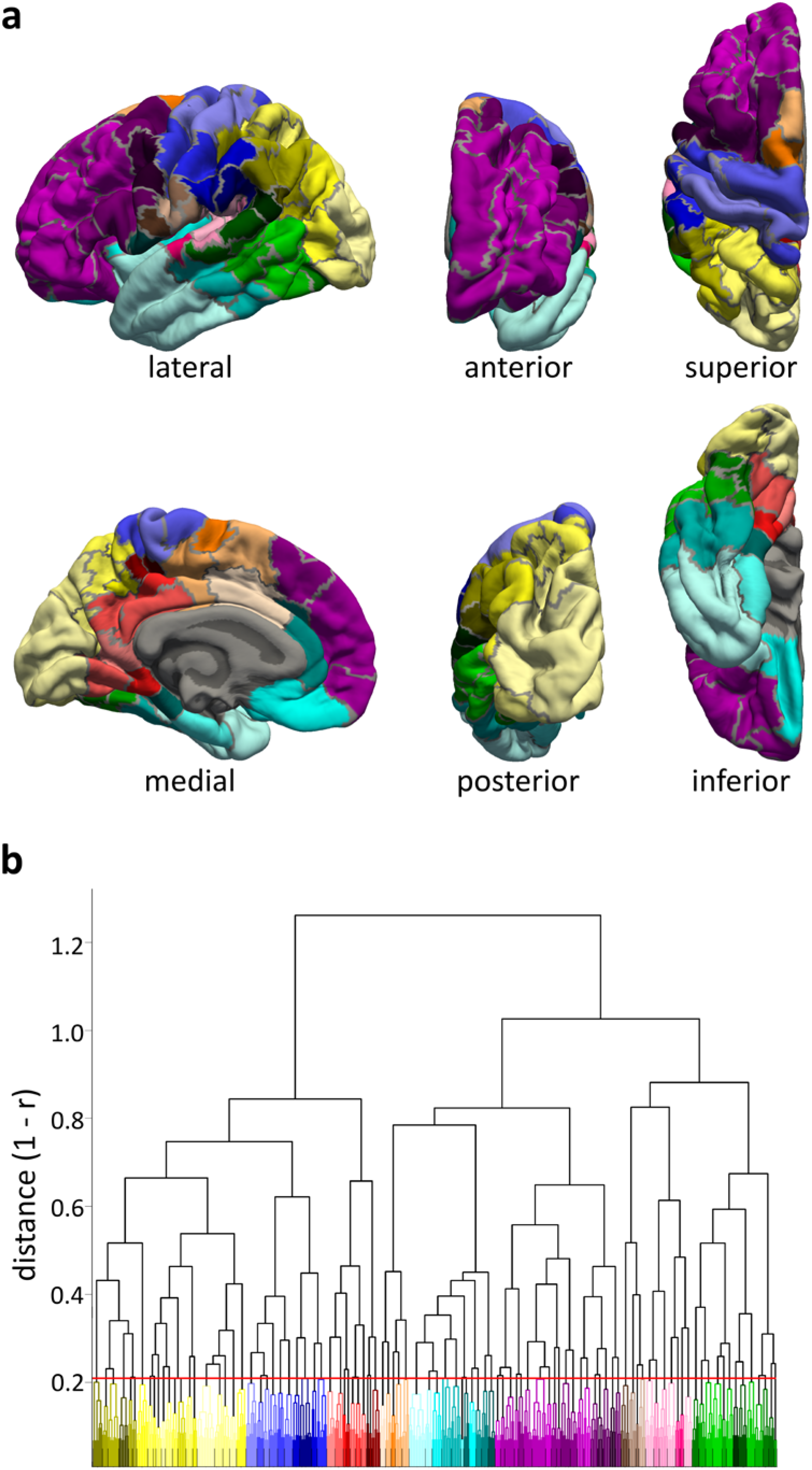
Transcriptome based parcellation of the human cerebral cortex. **a**) The parcellation into 100 clusters is displayed superimposed on the pial surface of the left cortical hemisphere of the fsaverage subject. **b**) The dendrogram created by the hierarchical clustering algorithm is shown with correlation distance calculated from Pearson’s r plotted on the y-axis. To reflect the structure of the dendrogram, which branches into 3 x 3 clusters at the highest levels, nine basic colors were used for display purposes. Further subdivisions down to the solution of 100 clusters are visualized by differences in brightness, darker clusters being more distant with respect to gene expression from the center of their respective higher-order cluster indicated by color assignment.

Inspecting the hierarchical relationships of cortical areas in the dendrogram from the top (Figure 1b), the first branching yields a posterior-dorsal cluster, containing the occipital and parietal lobes as well as the primary motor and posterior cingulate cortices (PCC), as well as an anterior-ventral cluster consisting of the frontal, temporal and mid- and anterior cingulate cortices (ACC). Subsequently, a third lateral cluster in the posterior temporal lobe is separated from the anterior-ventral cluster. In the next steps, the posterior-dorsal cluster is split into three clusters including the PCC and parts of the lingual gyrus (colored in red), the pre-, postcentral and supramarginal gyri corresponding to the primary motor, primary and secondary somatosensory areas (blue), and the parietal and occipital lobes (yellow), respectively. The anterior-ventral cluster is split into three clusters covering the majority of the frontal lobe (purple), most of the temporal lobe, insula, ACC and ventro-medial prefrontal cortex (cyan), and the mid-clingulate cortex and posterior parts of the superior frontal gyrus (orange). The lateral cluster is split into three clusters including parts of the superior temporal gyrus, temporal and parietal opercula covering most of the auditory cortex (pink), the majority of the posterior temporal lobe (green) and a lateral fraction of the pre- and post-central gyri (brown).

### Gene set enrichment analysis (GSEA)

GSEA was applied to the transcriptomic profiles of each cluster to test if disease-associated gene sets were over-represented among high expressing genes. GSEA identified significant (false discovery rate (FDR) controlled at q≤0.05) positive enrichment signals in at least one cortical cluster for 43 GWAS-derived and 84 expert curated trait-associated gene sets. A network based on the overlap between these gene sets is shown in Figure 2. Notably, no connections between GWAS and curated gene sets are depicted, because only edges with an overlap coefficient above 0.5 are included. The highest overlap coefficients between gene sets of different sources are between bipolar disorder (GWAS) and manic disorder (curated) and depression, bipolar (curated): overlap coefficient=0.38 and 0.35, respectively. Figures displaying enrichment patterns for all gene sets with significant signal in at least one cluster are available on our homepage. The Supplementary Table contains a list of all 266 tested gene sets, GSEA results for all significant enrichments and the results of leading edge analysis including an expression-weighted score reflecting the contribution of each gene to the enrichment of a gene set across all its significant clusters.

**Figure 2.**
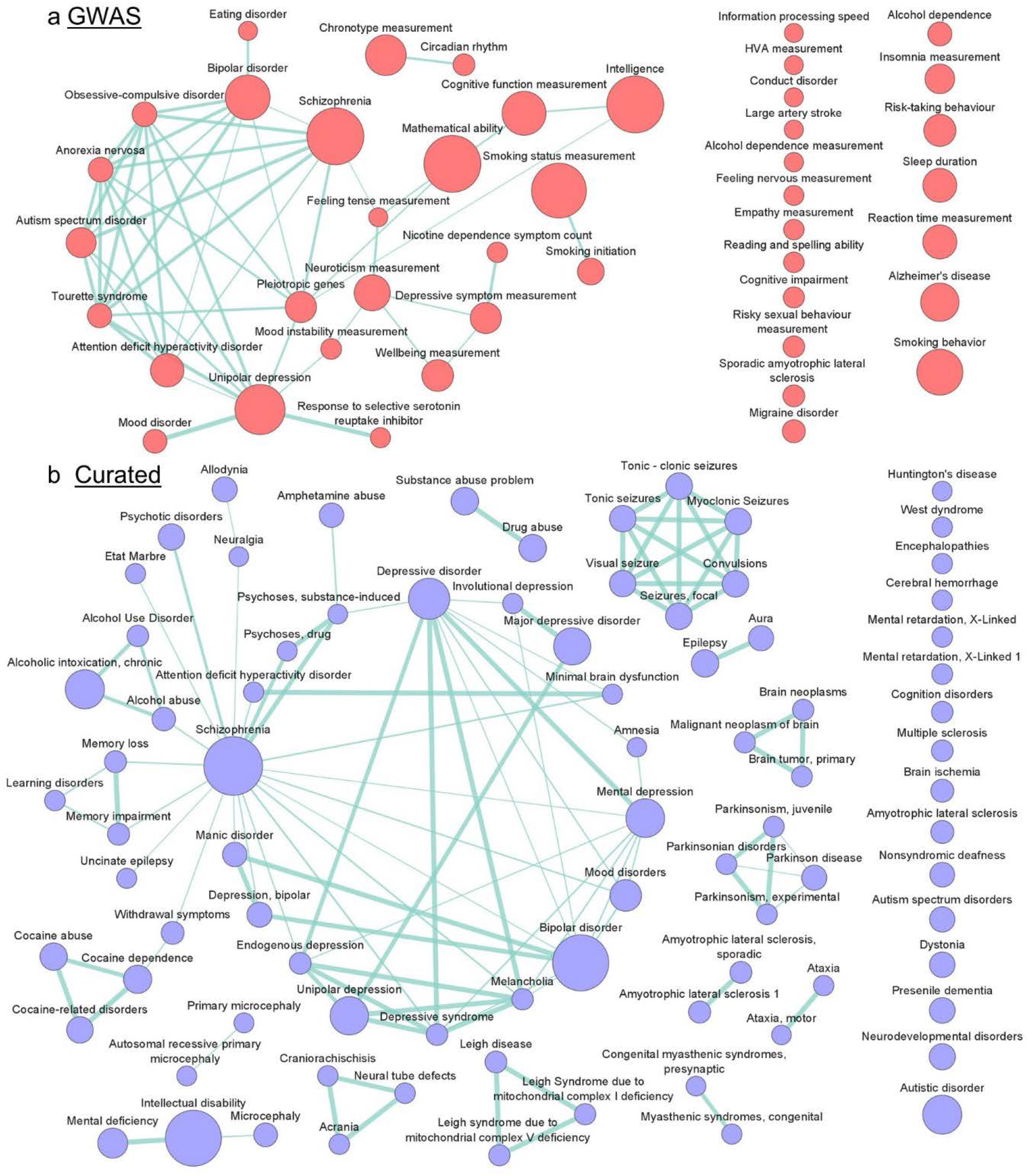
Network of gene sets enriched in cortical areas delineated with transcriptome-based parcellation. Gene sets derived from **a**) GWAS and **b**) curated gene-disease associations with significant positive enrichment in at least one cortical cluster are shown. Sizes of nodes are proportional to gene set sizes. Edges are included if the overlap coefficient for two gene sets is at least 0.5.

In the following paragraphs, we summarize the main findings of GSEA for principal disease groups. The discussed traits and their highest-scoring genes are listed in Table 1.

**Table 1.**
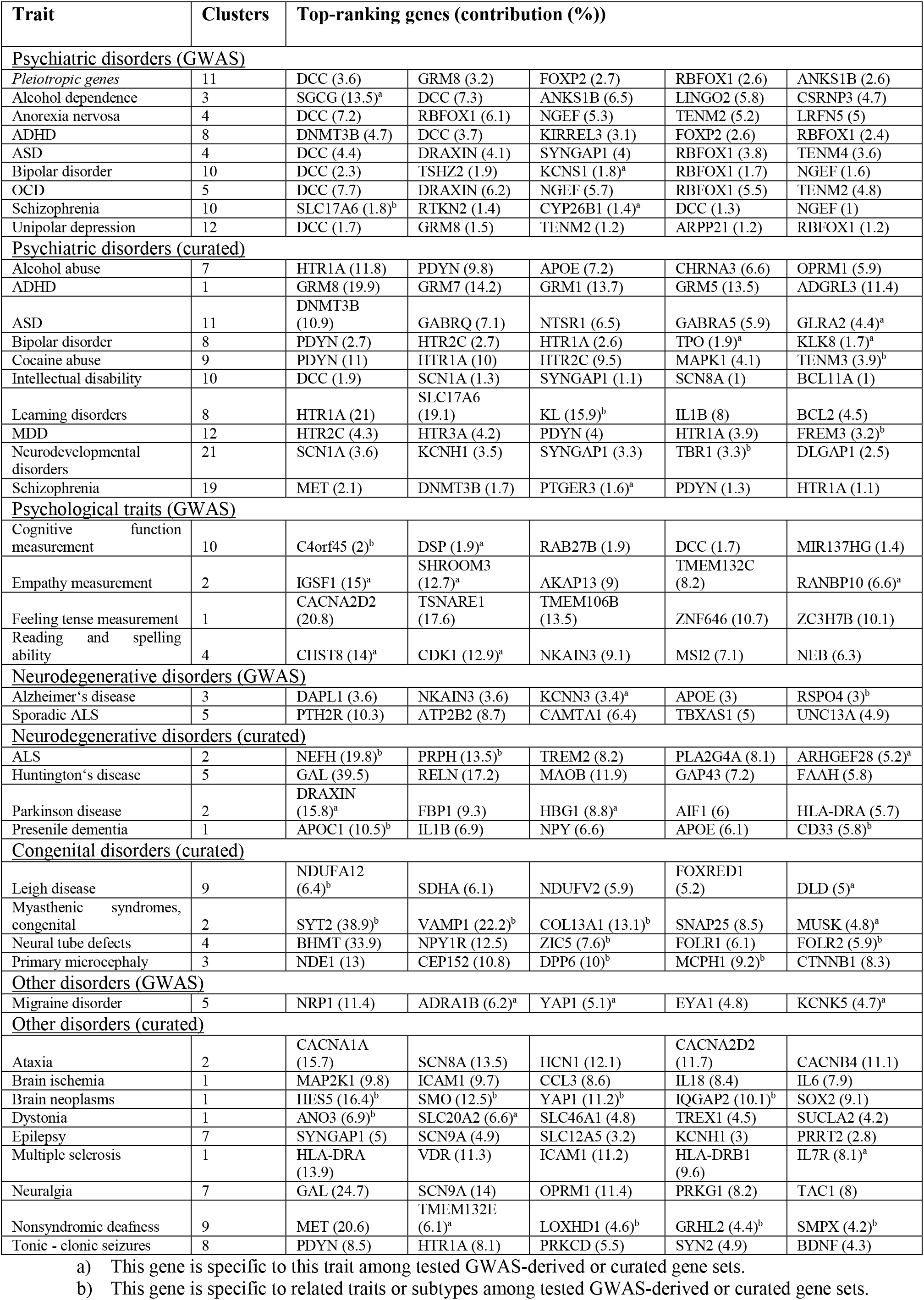
Genes with the highest contribution to enrichment signals of trait-associated gene sets. We calculated the relative contribution of each gene to the enrichment signal of its gene set based on its expression in significant clusters. This table lists the five top-ranking genes for each gene set presented in the result section. Scores for all gene sets and genes are provided in the Supplementary Table. The column “clusters” specifies the number of clusters with significant enrichment of a trait’s gene set. ADHD, attention deficit hyperactivity disorder; ALS, amyotrophic lateral sclerosis; ASD, autism spectrum disorders; MDD, major depressive disorder; OCD, obsessive-compulsive disorder.

### Psychiatric disorders

We observed the majority of significant enrichment signals for gene sets of psychiatric disorders. Enrichment patterns for common psychiatric disorders are displayed in Figure 3.

**Figure 3.**
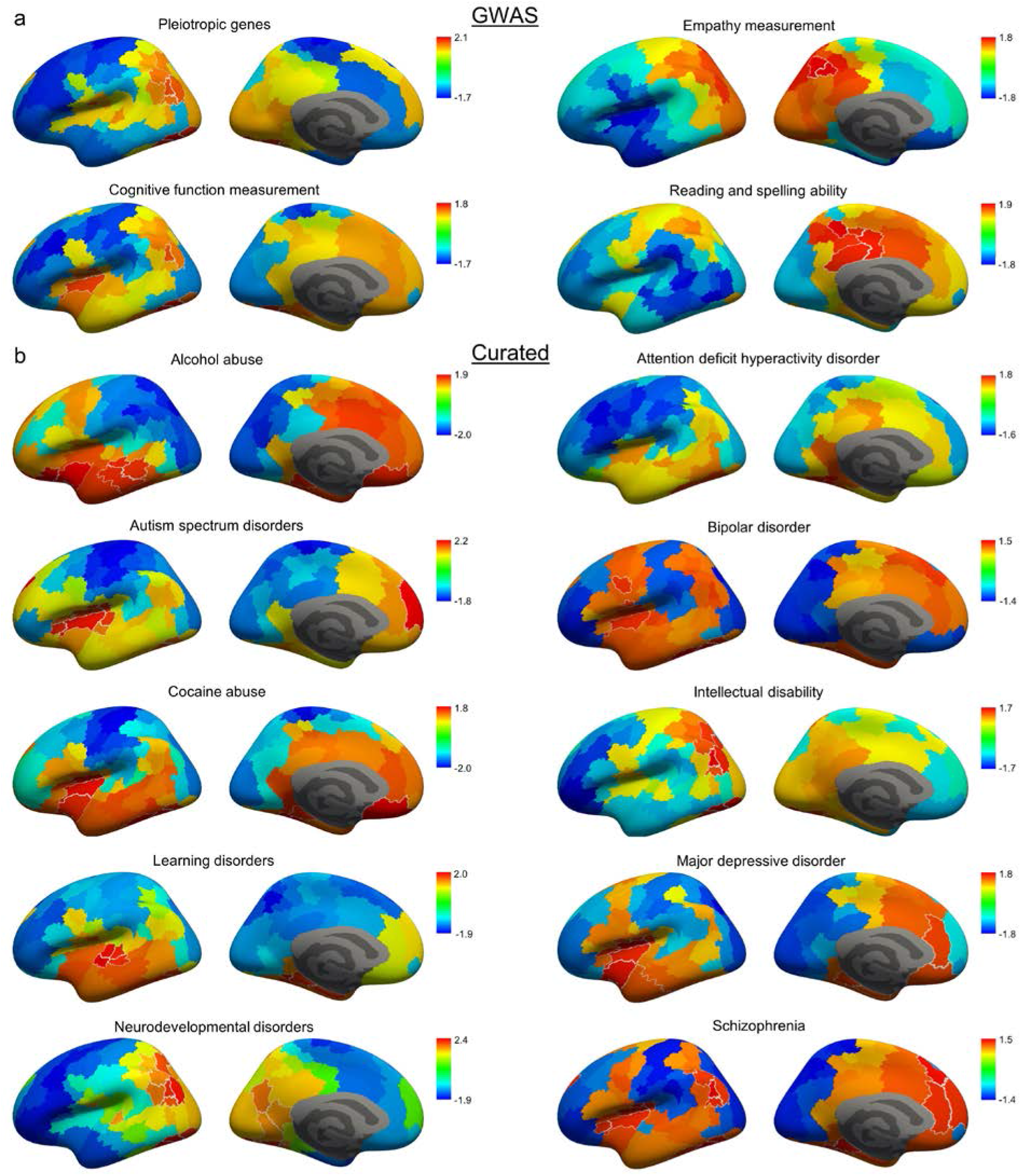
Cortical enrichment patterns of gene sets associated with psychiatric disorders and psychological traits. Normalized enrichment scores (NES) for each cortical cluster are displayed on an inflated representation of the cortical surface. **a**) The enrichment pattern of a set of pleiotropic genes associated with at least ten brain disorders by GWAS is shown, as it is highly similar to the patterns observed for GWAS-derived gene sets for individual psychiatric disorders. However, GWAS-derived genes associated with psychological traits and measurements had distinct enrichment patterns. **b**) Enrichment patterns of curated gene-sets for common psychiatric disorders are shown. White borders highlight significant clusters (FDR q≤0.05).

A high similarity of enrichment patterns of GWAS-derived gene sets was evident with significant enrichment in the inferior parietal lobule and posterior fusiform gyrus for most traits. This is in line with the high overlap coefficients between gene sets visible in Figure 2b. Based on this, we created a gene set containing genes mapped to a minimum of 10 brain disorders or psychological traits via GWAS. This set of pleiotropic genes was enriched in the same areas (Figure 3b). In accordance, genes from the pleiotropic set were among the ten highest-ranking for each psychiatric disorder. Only *SGCG* (14.0%), which was specific to alcohol dependence, i.e. not in any other GWAS-derived gene set analyzed, had a markedly higher contribution to enrichment signal compared to genes within the pleiotropic set. Other high-ranking genes specifically associated with traits were *KCNS1* (1.8%) for bipolar disorder and *CYP26B1* (1.4%) for schizophrenia. Of note, the relative contribution score of individual genes must be considered in the context of the large size of these gene sets.

Contrary to GWAS-derived gene sets, enrichment patterns for expert curated gene sets for psychiatric diagnoses were more diverse (Figure 3b). In line with this, despite significant overlaps between gene sets, a clear contrast in connection strengths within and across diagnostic subgroups was discernible (Figure 2b). In short, gene sets related to depressive disorders were enriched in insula, ACC, entorhinal cortex, parahippocampal gyrus and the rostral temporal cortex, which was driven by the expression of serotonin receptors, but also by *PDYN* and *FREM3* (3.2% for major depressive disorder (MDD)) with the latter being specifically associated with depression. Bipolar disorder showed enrichment in the posterior fusiform gyrus, insula and dorsolateral prefrontal cortex with a high score for specifically associated genes *TPO* (1.9%) and *KLK8* (1.7%) next to serotonin receptors. Curated gene sets associated with neurodevelopmental disorders and intellectual disability had similar enrichment patterns to the GWAS-derived set of pleiotropic genes. Strongly contributing genes were also associated with schizophrenia, autism and epilepsy. Genes associated with autism spectrum disorders (ASD) were enriched in the insula, mPFC and posterior fusiform gyrus with a high contribution of *DNMT3B* (10.9%), gamma-aminobutyric acid receptors and the specifically associated *GLRA2* (4.4%). Genes associated with learning disorders were enriched in the entorhinal cortex, parahippocampal, superior frontal and posterior fusiform gyri, which is similar to the patterns of the overlapping gene sets for memory loss and – impairment. The klotho gene, *KL*, is associated with these three traits and strongly contributes to their enrichment. The curated gene set for attention deficit hyperactivity disorder (ADHD) was enriched in a single cluster in the posterior fusiform gyrus with a strong contribution from genes coding for metabotropic glutamate receptors. Schizophrenia was enriched in the posterior fusiform and parahippocampal gyri, the inferior parietal lobule, as well as in the insula and medial prefrontal cortex (mPFC) with contribution of the specifically associated gene *PTGER3* (1.6%). The enrichment signal in the insula overlapped with those of drug induced psychoses and amphetamine abuse. Further, genes associated with alcohol and cocaine abuse were enriched in the insula, medial oribitofrontal cortex and parahippocampal gyrus with *TENM3* (3.9%) being specifically associated with cocaine-related traits.

### Psychological traits and measurements

In contrast to GWAS-derived gene sets for psychiatric diagnoses, different enrichment patterns were observed for behavioral traits and measurements not allocated to specific diseases. The traits “reading and spelling ability” and “empathy measurement” had the most distinct patterns with enrichment in the PCC and precuneus (Figure 3a) driven by expression of *CHST8* (14.0%) and *CDK1* (12.9%), and *IGSF1* (15.0%) and *SHROOM3* (12.7%) specific to these traits, respectively. “Cognitive function measurement” was enriched in the same areas as the pleiotropic set, but had additional signal in the area of the posterior insula with *TFAP2D* being specific and the highest expressing gene of the leading edge of the set in this area.

### Neurodegenerative disorders

Enrichment patterns of neurodegenerative disorders are displayed in Figure 4. The GWAS-derived gene set for Alzheimer’s disease (AD) had enrichment signals in the mPFC and entorhinal cortex with high signal from *DAPL1* (3.6%), *KCNN3* (3.4%) and *RSPO4* (3.0%). Similarly, the curated gene set for presenile dementia was enriched in the entorhinal cortex with strong signal from *APOC1* (10.5%) and *CD33* (5.8%), genes it shared exclusively with the curated gene set for AD. The curated gene set for Parkinson’s disease (PD) was enriched in the entorhinal cortex and frontal operculum with specific signals from *DRAXIN* (15.8%) and *HBG1* (8.8%). However, while *DRAXIN* was specific to PD in the curated data, it was also associated with ASD and OCD via GWAS. The curated gene set for Huntington’s disease was enriched in the mPFC and superior frontal gyrus with a high contribution of *GAL* (39.5%) and *RELN* (17.2%). The curated gene sets for amyotrophic lateral sclerosis (ALS), ALS 1 and sporadic ALS were enriched in the precentral gyrus with high contribution from the specifically associated genes *NEFH* (19.9%), *PRPH* (13.5%), *ARHGEF28* (5.2%), while the GWAS-derived gene set for sporadic ALS showed enrichment in the superior and inferior parietal cortex.

**Figure 4.**
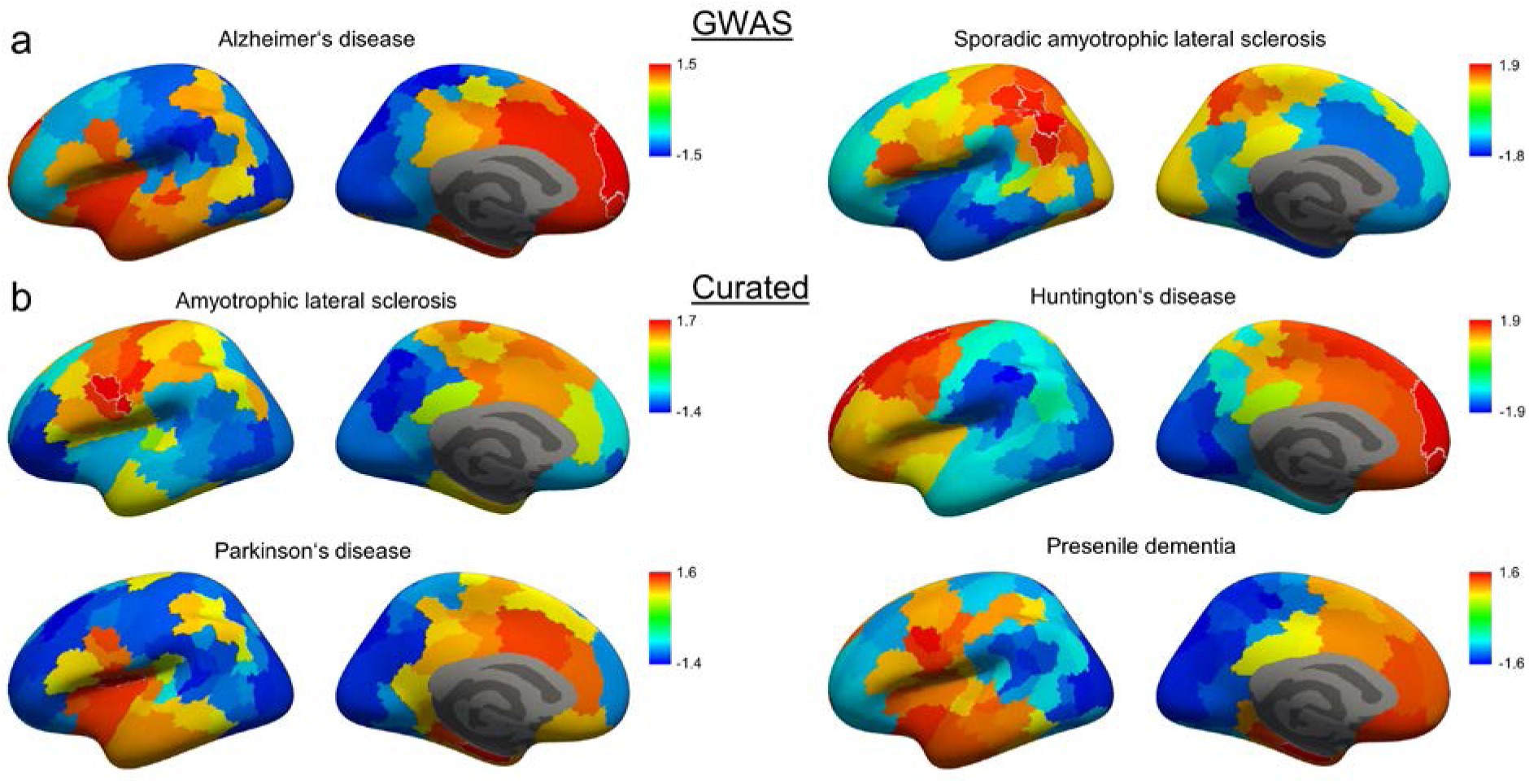
Cortical enrichment patterns of gene sets associated with neurodegenerative disorders. Normalized enrichment scores (NES) for each cortical cluster are displayed on an inflated representation of the cortical surface. **a**) Enrichment patterns of GWAS-derived and **b**) curated gene sets are shown. White borders highlight significant clusters (FDR q≤0.05).

### Congenital disorders

Significant enrichment signals for several curated gene sets for congenital disorders were observed (Figure 5). The enrichment of genes associated with congenital myasthenic syndromes was strongly driven by *SYT2* (38.9%), *VAMP1* (22.2%) *COL13A1* (13.1%) and *MUSK* (4.8%), which were specific to myasthenic syndromes. Genes associated with Leigh disease were enriched in the pre- and postcentral gyri and superior parietal cortex with specific signal from *NDUFA12* (6.4%) and *DLD* (5.0%). The gene set associated with neural tube defects was enriched around the frontal and temporal poles with high contribution from *BHMT* (33.9%) and specific signal from *ZIC5* (7.6%), *FOLR2* (5.9%) and *VANGL2* (4.9%), which it shared with the highly overlapping gene sets of acrania and craniorachischisis. Genes associated with primary microcephaly were enriched in clusters in the lateral occipital cortex and superior temporal gyrus with specific signal from *DPP6* (10%) and *MCPH1* (9.2%).

**Figure 5.**
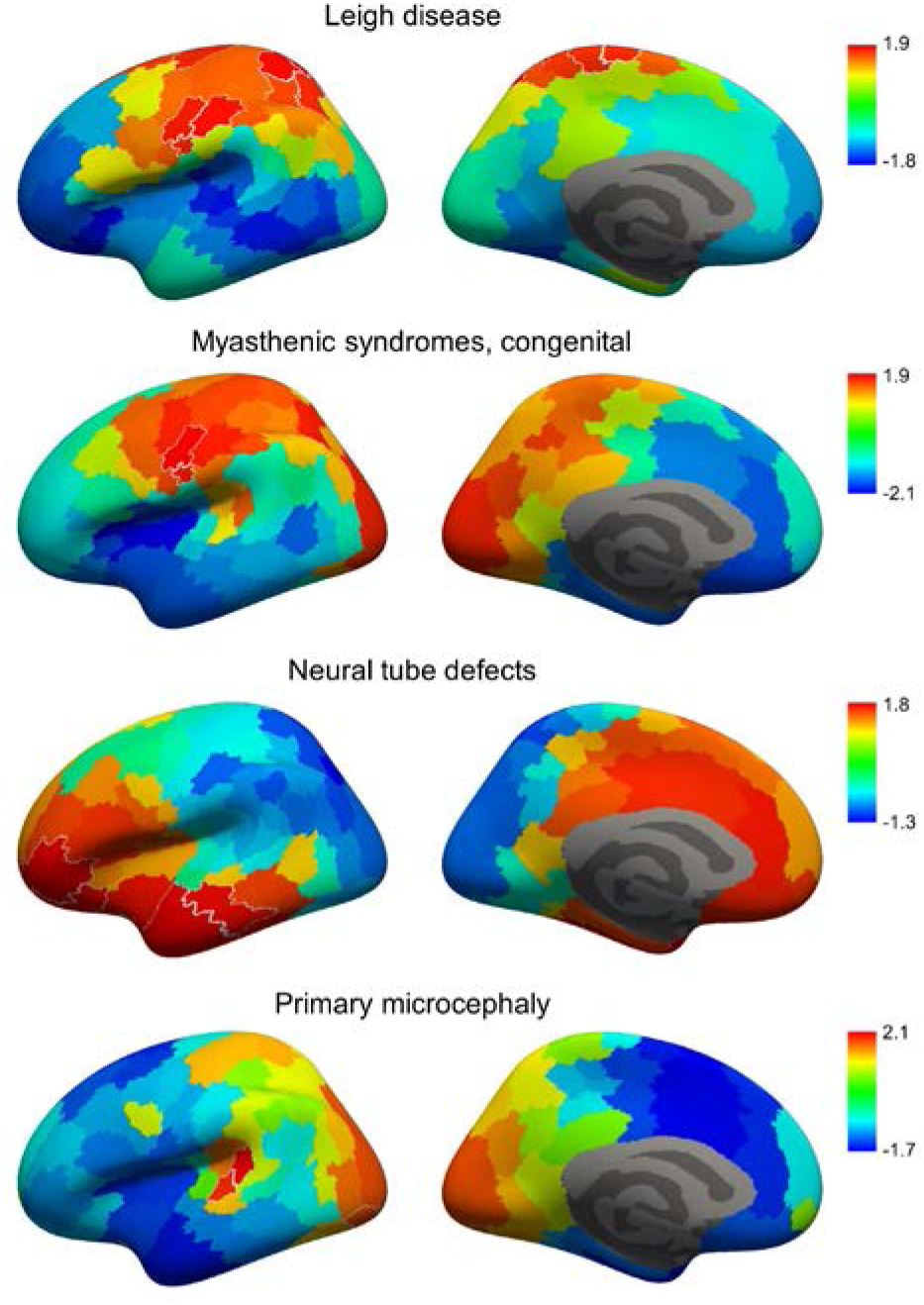
Cortical enrichment patterns of gene sets associated with congenital disorders. Normalized enrichment scores (NES) for each cortical cluster are displayed on an inflated representation of the cortical surface. White borders highlight significant clusters (FDR q≤0.05).

### Other neurological disorders

The curated gene set for epilepsy was enriched in the posterior fusiform gyrus and lateral occipital cortex, while the gene set for tonic-clonic seizures, which is highly overlapping with the gene sets of other forms of seizures, was enriched in the mPFC and the middle part of the inferior, middle and superior temporal and posterior fusiform gyri (Figure 6). Genes associated with ataxia were enriched in the superior parietal lobule and lateral occipital cortex with the majority of the signal coming from genes coding for ion channels. *ANO3* (6.9%) and *SLC20A2* (6.6%) were specifically associated with dystonia and were the strongest contributors to the enrichment signal in the medial precentral gyrus. Genes coding for proteins involved in immune responses strongly contributed to the enrichment signal of brain ischemia in the frontal operculum *(MAP2K1, ICAM1, CCL3, IL18* and *IL6*) and multiple sclerosis in the inferior frontal gyrus *(HLA-DRA, VDR*, *ICAM1, HLA-DRB1, IL7R).* Genes associated with brain neoplasms were enriched in the mid-cingulate cortex with several specific genes contributing to the signal. The curated gene set associated with neuralgia and the GWAS-derived gene set for migraine disorder were both enriched in the mid-cingulate cortex, while only *PRKG1* was shared between the two sets. Several specific genes contributed to the enrichment of migraine disorder: *ADRA1B* (6.2%), *YAP1* (5.1%) and *KCNK5* (4.7%). Lastly, the curated gene set for nonsyndromic deafness was enriched in parts of the middle- and superior temporal gyri, as well as in the PCC and lingual gyrus with a number of specific genes contributing to its signal: *TMEM132E* (6.1%), *LOXHD1* (4.6%), *GRHL2* (4.4%) and *SMPX* (4.2%).

**Figure 6.**
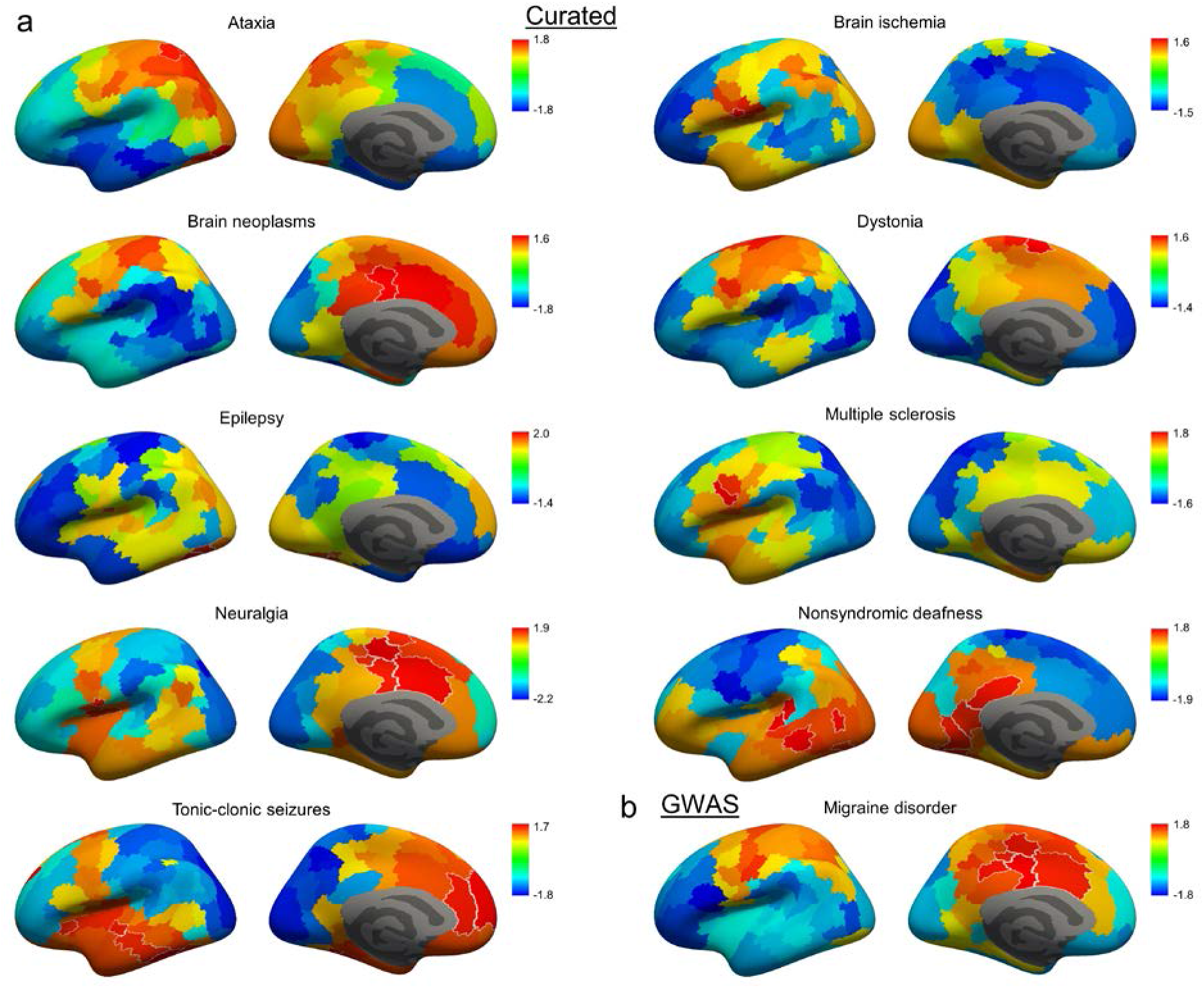
Cortical enrichment patterns of gene sets associated with miscellaneous brain disorders. Normalized enrichment scores (NES) for each cortical cluster are displayed on an inflated representation of the cortical surface. **a**) Enrichment patterns of curated and **b**) GWAS-derived gene sets are shown. White borders highlight significant clusters (FDR q≤0.05).

## Discussion

We have developed and demonstrated the utility of a data analysis paradigm for the integration of transcriptomic data with gene-disease association databases, which allows for the creation of novel insights into the functions and disorders of the human cerebral cortex. Firstly, the transcriptome-based parcellation allows for an efficient assessment of the similarity in gene expression profiles between cortical areas in the adult human brain. Solely based on transcriptomic data, hierarchical clustering was able to delineate borders in line with major anatomical landmarks, as well as with functional differentiation of primary motor, somatosensory, visual and auditory cortices. While this is beyond the scope of this work, it will be interesting to see how the delineated areas evolve across developmental stages.^15,16^

The continuous expression data enabled the extraction of unbiased, normalized and distinct transcriptomic profiles from transcriptome-based clustering for GSEA. Gene-disease associations derived from curated databases differ from GWAS results as they include information based on targets for treatment and toxins, disease models and molecular pathways among other sources. We show that genes with pleiotropic effects in GWAS studies are strongly enriched in the posterior fusiform gyrus (FG) and inferior parietal lobule (IPL). Our expression-based score identified *DCC* and *RBFOX1* as leading contributors to this signal. This is in line with a recent meta-analysis of genome wide studies which found variants associated with these genes to have pleiotropic effects across multiple psychiatric disorders, the involvement of pleiotropic loci in neurodevelopmental pathways and their enrichment among genes with high pre- and postnatal expression in neuronal cell types.^17^ The relevance of the enrichment of these genes in these specific regions of the adult brain needs to be established. The fusiform gyrus is a hominoid-specific structure and the role of the posterior FG in face recognition is well-established.^18^ Remarkably, the allocation of the posterior FG and the IPL to same cluster based on multi-receptor data derived from autoradiography of adult humans implicated a common functioning of these regions, especially of the caudal portion of the IPL.^19^ While IPL lesions are associated with hemi-neglect,^20^ functional imaging has revealed its role in semantic processing serving social cognition and language.^21,22^ Alterations in response to faces and social cues have been documented in most psychiatric disorders with meta-analytic evidence for different activation and structural alterations in the IPL and FG for patients suffering from schizophrenia,^23,24^ autism, ^25,26^ ADHD,^27^ and depression.^28–30^ In line, reductions in neuron density in the FG of patients diagnosed with schizophrenia^31^ and autism^32^ were reported. The alignment of the enrichment pattern of genes with pleiotropic effects in GWAS with that of the curated gene set associated with neurodevelopmental disorders supports the notion that circuit formation in these areas during critical periods might affect the unspecific predisposition to develop a mental disorder later in life. In contrast to psychiatric disorders, we observed distinct enrichment patterns for genes associated with a number of psychological traits via GWAS. This might lend support to a dimensional approach for the study of the genetic basis of brain functions. The enrichment of genes specifically associated with empathy in the precuneus area is remarkable given the replicated findings of activation in the area in tasks involving mentalization.^33^

The enrichment of genes associated with the movement disorders ALS, myasthenic syndromes and dystonia in the motor cortex, and genes associated with memory loss, Alzheimer’s disease and presenile dementia in the entorhinal cortex lends support to our method based on the well-established functions of these cortical areas. The potential that enrichment analysis might suggest hitherto unknown roles of cortical areas in brain disorders may constitute the first step towards the discovery of novel disease mechanisms and treatments through experimental and clinical validation. We will explore some hypotheses inspired by the results in the following sentences. The enrichment of genes associated with Huntington’s disease in the frontal cortex is in line with cognitive and psychiatric symptoms and implies that local processes beyond the degeneration of striato-cortical projections contribute to the neuronal loss observed in this area.^34^ The enrichment signal in the entorhinal cortex and frontal operculum of genes associated with Parkinson’s disease might correspond to the development of pseudobulbar symptoms and cognitive impairments in some cases.^35^ The high contribution of genes coding for proteins involved in immune signaling to the enrichment signal of brain ischemia in the frontal operculum suggests that a dysregulated immune response might underlie the vulnerability of this area to hypoxia, as seen in bilateral opercular syndrome.^36,37^ The enrichment of genes associated with multiple sclerosis in the inferior frontal gyrus might explain reductions in blood flow and activation correlated with cognitive impairments in affected patients.^38,39^ Lastly, the enrichment of genes associated with neuralgia and migraine in the midcingulate cortex is in accordance with a recently characterized pathway from the area to the posterior insula that supports the development of chronic hypersensitivity to pain.^40^

Limitations arising from different sources need to be considered. Firstly, transcriptomic data used was obtained from the left hemisphere of only six healthy adult individuals. Given the macrodissection of samples, different cell types from different cortical layers were analyzed. The registration of brain samples to magnetic resonance imaging data acquired post-mortem and further to a standard representation of the cortical surface might have introduced distortions. We had to merge the sparse data of these individuals in order to obtain a single estimate of cortical gene expression. The isotropic prediction model and the smoothing necessary to reduce the influence of outliers might have further blurred the borders between brain areas. GSEA results are critically depending on careful curation of the gene-disease association databases, which are refined continuously. Lastly, our analysis of GWAS-derived gene sets was limited to variants localized within genes, which provides a narrow representation of the entire basis of heritability.

In summary, the combination of transcriptome-based parcellation of the cerebral cortex with gene set enrichment analysis is a powerful method to gain novel insights into the involvement of cortical areas in brain functions and disorders. The identified enrichments of disease-associated genes suggest the vulnerability of specific cortical areas to noxae of various origins, which might alter the risk to develop one or several brain disorders. For several diseases, specific genes were highlighted which could be prioritized in the development of novel disease models and treatments.

## Methods

### Estimation of comprehensive cortical gene expression

Human brain transcriptome data provided by the Allen Institute (Allen Human Brain Atlas, http://human.brain-map.org/) was downloaded and processed as described previously^13^ to generate comprehensive maps of gene expression in the cortex which are openly available on our homepage (http://www.meduniwien.ac.at/neuroimaging/mRNA.html). In short, microarray probes with signal not significantly different from background in a minimum of 1% of samples and probes without an association with an Entrez Gene ID were excluded, such that 40,205 probes mapped to 18,686 genes were retained. Probes associated with the same gene were averaged and individual probes were removed step-wise if their exclusion resulted in a higher spatial dependence of gene expression. For each gene, expression intensity in the six donor brains was set to an equal mean across brain regions to minimize the influence of inter-individual variation. Cortical surface reconstruction was performed using the recon-all pipeline in FreeSurfer 5.1 (Harvard Medical School, Boston, USA; http://surfer.nmr.mgh.harvard.edu/) and microarray samples from all donors were mapped to their respective nearest vertices in fsaverage space. Variogram models were fitted and Gaussian process regression (ordinary Kriging) was performed to predict mRNA expression for all surface vertices using the gstat 1.1-5 package^41^ in R. This was performed for the left hemisphere, as data for the right hemisphere was not available for 4 out of 6 brains. In order to reduce noise, all original samples were replaced by the average of the estimated expression intensity of directly adjacent vertices. Then, smoothing was applied with an isotropic Gaussian kernel corresponding to four edges in fsaverage space (approximately 3.4 mm full width at half maximum). Normalization of each gene expression map was achieved by subtraction of the mean expression intensity across the cortex.

### Parcellation of the cortex

18,686 gene expression maps were used as input for agglomerative hierarchical clustering in Matlab R2018a (https://www.mathworks.com/). The distance between two vertices of the cortex was calculated from Pearson correlation coefficients (r) using the formula d(x,y)=1−r_x,y_, which provides high accuracy while being computationally efficient.^42^ A correlation based distance was used, because the contrast between nearest and furthest neighbors decreases with an increasing number of dimensions for Minkowski measures, such as Euclidean or Manhattan distances, which are otherwise frequently used in clustering applications.^43^ Robustness analysis was performed in order to select the optimal linkage method (average, complete or median linkage) and assess effect of scaling gene expression data to the unit standard deviation. Scaling was predicted to reduce robustness due to the assignment of more weight to genes with a low spatial variance in expression. Robustness analysis was performed by creating 1,000 random subsets of 6,229 genes, i.e. one third of the entire transcriptome dataset, and performing clustering analysis on each of these subsets. Subsequently, the average of the Adjusted Rand Index (ARI)^44^ obtained for each pair of clustering results obtained from the 1,000 subsets and each number of clusters from k=5 to k=100. The ARI is a measure ranging between −1 and 1 which reflects the concordance between different partitions. In order to achieve computational feasibility, robustness analysis was performed on a downsampled representation of the cerebral cortex (fsaverage5) comprising 10,242 vertices. Results of robustness analysis are displayed in Supplementary Figure S2 and indicated highest robustness for average linkage, which is in accordance with a published analysis of the optimal choice of linkage method for clustering of gene expression data.^42^ As predicted, scaling to the standard deviation reduced robustness of clustering results irrespective of linkage method. On this basis, agglomerative hierarchical clustering with Pearson correlation distance, average linkage and without scaling was used as the final clustering algorithm.

Transcriptome-based parcellation was compared to the following established parcellations: cytoarchitectonic parcellation into Brodmann Areas (BA),^45^ clustering using resting-state functional MRI data into 7 networks by Yeo and collegues,^46^ and into 200 parcels per hemisphere using gradient-weighted Markov Random Field models by Schaefer and collegues,^47^ clustering based on structural connectivity assessed using diffusion-weighted MRI (Brainnetome Atlas),^48^ parcellation based on macroanatomical landmarks (Desikan-Killiany Atlas^49^), clustering based on the heritability of cortical thickness and surface area by Chen and collegues,^50^ and parcellation based on multiple MRI modalities by Glasser and collegues.^51^ For each comparison, the number of transcriptome-based clusters was set equal to the number of regions defined by the respective published parcellation. The overall agreement between different atlases was quantified using ARI.^44^ The similarity between each pair of regions was quantified using Dice’s coefficient as displayed in Supplementary Figure S1.^52^

### Gene set enrichment analysis

Gene set enrichment analysis (GSEA)^53^ was carried out in order to assess if gene sets associated with brain diseases and related phenotypes were overrepresented among highly expressing genes of cortical areas defined by transcriptome-based parcellation.

Gene sets were created based on gene-trait associations in the NHGRI-EBI GWAS Catalog v1.0.2^54^ and DisGeNET database^14^ which were downloaded in May 2020. Expert curated associations were extracted from the DisGeNET database for traits identified by a Unified Medical Language System^®^ Concept Unique Identifier (UMLS CUI) assigned to the MeSH classes C10 (nervous system diseases), F01 (behavior and behavior mechanisms) or F03 (mental disorders). In order to obtain gene-trait associations from the GWAS Catalog, SNPs located within genes and associated with Experimental Factor Ontology (EFO) traits were extracted. EFO traits were included if they were a subclass of nervous system diseases, psychiatric disorders or mental processes. Several traits had to be removed or added manually, e.g. to exclude eye disorders, myopathies or to include responses to psychopharmacological drugs. An additional gene set comprising pleiotropic genes associated with a minimum of ten different traits was created based on GWAS data. Gene sets with less than 15 genes were excluded as these might yield less robust results.^53^ Duplicated gene sets were removed. These steps resulted in a total of 266 gene sets (Supplementary Table).

The median of normalized expression for each gene and cortical parcel *k* was extracted. GSEA was performed using R 4.0.1 and the fgsea package 1.14.0.^55,56^ For each gene set *s* and *k*, 10,000 gene set permutations were performed in order to calculate nominal p-values and normalized enrichment scores (NES). Normalization is performed to the mean enrichment score of random gene sets of the same size. Analysis was performed at *k*=100, as it allowed for the assessment of enrichment signal at a resolution which enables appropriately precise anatomical localization with high robustness of clustering (Supplementary Figure S2) and expected high within-parcel homogeneity. False-discovery rate (FDR) was controlled at q≤0.05 across all *s* and *k* with adjustment for the estimate of the proportion of true null hypotheses.^57^ Only positive enrichments were reported as significant. The EnrichmentMap Cytoscape App 3.3 was used to visualize relationships between significant genesets.^58^

Finally, the relative contribution of individual genes to the enrichment of each significant geneset was estimated. To this end, the leading edge (LE) genes were extracted for each parcel in which a significant positive enrichment of a gene was observed. The LE genes are those which account for the enrichment signal of a geneset, i.e. the enrichment score is lower at all genes with a lower expression than the LE. For each significant parcel, the contribution of each gene in the LE was calculated as its expression divided by the sum of the expression of all genes in the LE. The overall contribution of a gene across all significant parcels was calculated by weighting its contribution in each parcel by the NES of the geneset.

## Data availability

Parcellation results in FreeSurfer (.annot) file formats and expression data for each gene and cluster are available for download at http://www.meduniwien.ac.at/neuroimaging/mRNA.html.

## Acknowledgements

MM is funded by the Austrian Science Fund (FWF), DOC 33-B27. KM and RMB are recipients of a DOC fellowship of the Austrian Academy of Sciences at the Department of Psychiatry and Psychotherapy, Medical University of Vienna. This scientific project was performed with the support of the Medical Imaging Cluster of the Medical University of Vienna.

## Author contributions

GG carried out GSEA, interpretation of results and wrote the manuscript. MM performed hierarchical clustering and robustness analysis. MP, KM and RMB provided support in statistical analysis. UJ supported interpretation of the results. LR provided guidance at all steps. All authors revised and approved the final manuscript.

