## Supplementary figures for "Enrichment of disease-associated genes in cortical areas defined by transcriptome-based parcellation"

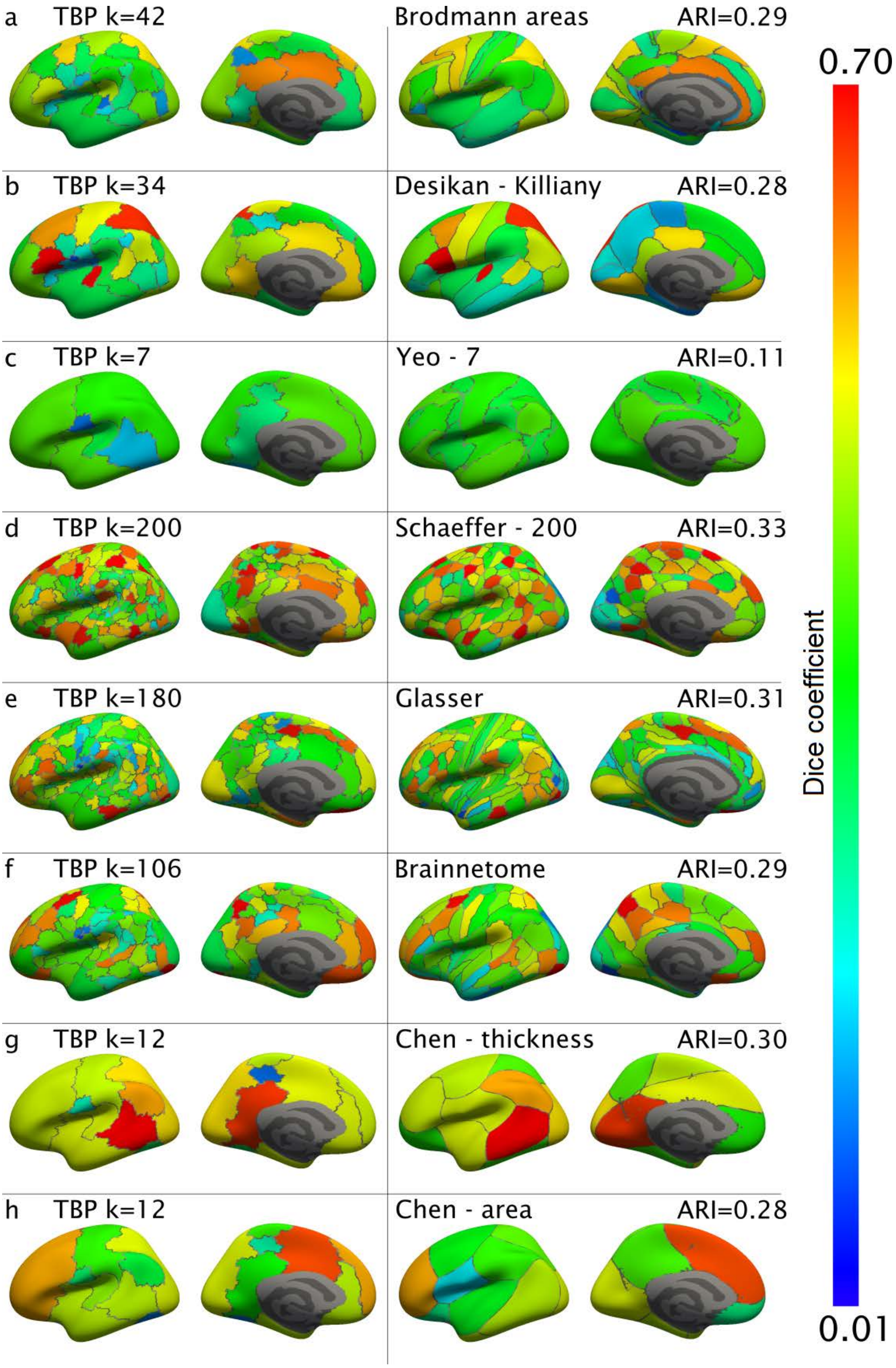

a
TBP k=42
Brodmann areas
ARI=0.29
b
TBP k=34
Desikan - Killiany
ARI=0.28
c
TBP k=7
Yeo - 7
ARI=0.11
d
TBP k=200
Schaeffer - 200
ARI=0.33
e
TBP k=180
Glasser
ARI=0.31
f
TBP k=106
Brainnetome
ARI=0.29
g
TBP k=12
Chen - thickness
ARI=0.30
h
TBP k=12
Chen - area
ARI=0.28
0.70
Dice coefficient
0.01

**Supplementary Figure S1: Comparison of transcriptome-based parcellation (TBP) with published parcellations based on different modalities.** The number of clusters was set equal in each comparison to facilitate interpretability. Each area was colored based on the highest Dice coefficient it shares with any area in the atlas which was compared, i.e. proportional to the overlap. Row by row, TBP is compared **a**) to Brodmann areas based on cytoarchitecture,[1] **b**) to the Desikan-Killiany Atlas based on macroanatomical landmarks, [2] **c**) to the parcellation into seven networks by Yeo and collegues based on resting-state functional MRI (rs-fMRI) data,[3] **d**) to parcellation into 200 areas using clustering based on rs-fMRI data using gradient-weighted Markov Random Field models as published by Schaefer and colleagues,[4] **e**) to the parcellation based on multiple MRI modalities (rs-fMRI, task fMRI, cortical thickness and myelination) by Glasser and colleagues,[5] **f**) to the parcellation based on structural connectivity assessed using diffusion-weighted MRI (Brainnetome Atlas),[6] and to parcellation using clustering based on the heritability of **g**) cortical thickness and **h)** surface area by Chen and collegues.[7] Results are displayed superimposed on an inflated representation of the left cortical surface. Despite the high overlap observed for certain areas in each atlas, overall, adjusted Rand index (ARI) ranged from 0.11 to 0.33. Notably, the maximum ARI observed between parcellations published by different authors was 0.37.

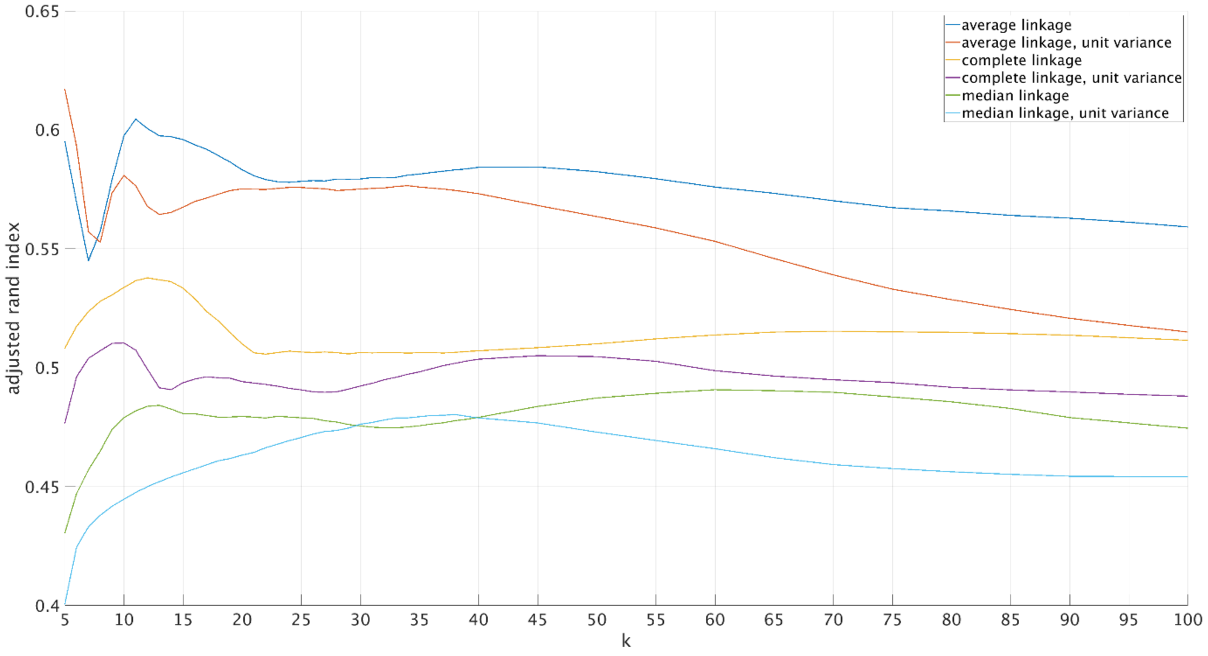


**Supplementary Figure S2: Analysis of robustness of hierarchical clustering methods of cortical gene expression data.** For each linkage method, average adjusted Rand Index (ARI) calculated for clustering results based on 1,000 random subsets comprising one third of the brain transcriptome dataset is plotted as a function of the number of clusters *k*. This was performed separately for normalized (each gene's expression set to zero mean) and scaled (z-scored) gene expression data. Highest robustness across a different range of *k* was observed for average linkage of normalized gene expression data. Scaling to the standard deviation reduced robustness of clustering results irrespective of linkage method and resulted in declining robustness with an increasing number of clusters for average linkage clustering.
